# The role of integrins in T cell-mediated resistance to *Cryptosporidium parvum*

**DOI:** 10.64898/2026.04.11.717894

**Authors:** Maria Merolle, Breanne E. Haskins, Julie Engiles, Andrew Hart, Ian S. Cohn, Christian Howard, Keenan O’Dea, Jessica Byerly, David Christian, Boris Striepen, Christopher A. Hunter

## Abstract

*Cryptosporidium* is a protozoan that infects epithelial cells of the small intestine and is a cause of diarrhea and death in immunocompromised individuals and malnourished children. Immunity to this parasite is mediated by an intestinal T cell response, which is generated in gut-associated lymphoid tissues and dependent on type 1 conventional dendritic cells (cDC1s). The initial priming of T cells is accompanied by changes in integrin expression and subsequent trafficking to the site of infection. The role of specific integrins in trafficking to the ileum during cryptosporidiosis is largely unknown. The development of a transgenic *Cryptosporidium* strain that expresses MHCI and MHCII-restricted model antigens provides the ability to track T cell responses to this parasite. Our studies in this system revealed marked changes in the integrin profile of parasite-specific T cells as they are activated and traffic to the gut, and demonstrate that cDC1s contribute to the expression of the integrins ⍺4, β7, β1, and ⍺L. Surprisingly, blockade of the canonical gut-homing integrin ⍺4β7 does not impact the ability of parasite-specific T cells to access the gut. However, blockade of integrin ⍺L decreases the parasite-specific T cell frequency at the site of infection and delays control of parasite burden. These datasets highlight an ⍺4β7-independent mechanism of T cell trafficking to the small intestine and indicate that ⍺L is an integrin required for T cell-mediated resistance to *Cryptosporidium*.

## Introduction

The ability of activated T cells to home to the gut is important to mediate protection against many different enteric pathogens (1–3), including *Cryptosporidium spp* (*4*). *Cryptosporidium* preferentially infects the epithelial cells of the small intestine and is an important cause of diarrheal disease in individuals with defects in T cell function (5, 6), and a significant cause of disease in immunocompetent individuals (7, 8). In order for the host to control this infection, effector CD4^+^ and CD8^+^ T cells must traffic to the site of infection, where their production of interferon-ɣ restricts parasite growth (9–11). Integrins, transmembrane proteins composed of dimerized ⍺ and β subunits, are a key family of homing receptors that mediate T cell migration to peripheral tissues by binding ligands expressed by the target tissue (12, 13). The events that lead to effector T cell responses in the gut begin with CD103^+^ DCs that sample antigen and migrate from the gut mucosa to the mesenteric lymph node (mLN) or Peyer’s Patches (PP) to present antigen to naïve T cells (14, 15). These DCs express high levels of retinol dehydrogenase (RDH) and retinaldehyde dehydrogenase (RALDH1/2), the key enzymes that metabolize vitamin A into retinoic acid (RA) (16, 17). The local production of RA in the mLN signals through the RA receptor in T cells which leads to expression of the integrin dimer ⍺4β7, also called Lamina Propria Adhesion Molecule-1 (LPAM-1) (18–20). Seminal lymphocyte trafficking studies identified ⍺4β7 as a significant contributor to homing to the intestine through their interactions with Mucosal Addressin Cellular Adhesion Molecule 1 (MAdCAM-1) on endothelial cells of the gut (21–23).

Studying the contributions of integrins to T cell responses is important to understand a variety of intestinal pathologies. The ability of T cells to traffic to the gut underlies the development of protective and pathological responses and there has been a long-standing interest in these pathways. Most studies that have investigated the function of ⍺4β7 on T cells have been performed at homeostasis or in models of oral tolerance or colitis (24–27), with varying time frames of trafficking that span hours (21, 28) to weeks (29). Others have highlighted that less is known about the contribution of ⍺4β7 during enteric infections (30). Other integrins, such as αEβ7, α4β1, and αLβ2 have also been implicated in T cell trafficking to the intestine (31–36) but the relevance of these mechanisms to the anti-*Cryptosporidium* T cell response remains unclear.

To better understand T cell responses that mediate resistance to *Cryptosporidium* infection, mouse-adapted *Cryptosporidium parvum* parasites (37) have been developed that express the MHC class-I and II-restricted peptide model antigens SIINFEKL and gp61-80, derived from ovalbumin and LCMV (Lymphocytic Choriomeningitis Virus), respectively (38, 39). These model antigens are encoded in the parasite gene *medle-2,* which is exported from the parasite into the host cell cytosol (40). These transgenic parasites, known as maCp-ova-gp61, allow the use of TCR transgenic CD8^+^ OT-I and CD4^+^ Smarta T cells as surrogates for the parasite-specific T cell response (41). Here, we utilize this system to evaluate integrin expression on T cells responding to *Cryptosporidium* during priming in the mLN and as effectors in the gut. Based on these data, blocking antibodies were used to assess the contribution of different integrins to trafficking of parasite-specific T cells to the site of infection. The use of this model system highlights an ⍺4β7-independent mechanism of T cell trafficking to the small intestine and reveals integrin ⍺L as critical to resistance to *Cryptosporidium*.

## Material and Methods

### Mice

C57BL/6 (stock no: 000664), Nur77-GFP reporter mice, (stock no: 016617), CD45.1 C57BL/6 mice (stock no: 002014), *Irf8+32*^-/-^ mice (stock no: 032744), OT-I mice (stock no:003831), Smarta CD45.1 mice (stock no: 030450) and IFN-γ^-/-^ (stock no: 002287) were purchased from Jackson Laboratories and then maintained in-house. Mice used in this study were males or females ranging from 5 to 12 weeks, and all mice were age- and sex-matched within individual experiments. All protocols for animal care were approved by the Institutional Animal Care and Use Committee of the University of Pennsylvania (protocol #805405 and #806292).

### Mouse infection and measurement of parasite burden

Transgenic parasites were generated and isolated as previously described (42, 43). The parasites utilized were a version of mouse-adapted version of *Cryptosporidium parvum* modified to express the MHC class I-restricted antigen SIINFEKL (an epitope in the protein ovalbumin) and the MHC class II-restricted antigen gp61-80 (an epitope in an LCMV glycoprotein) (44). This strain is referred to as maCp-ova-gp61. Mice were infected with 5×10^4^ oocysts by oral gavage. To quantify fecal oocyst shedding, 20 mg of pooled cage feces was suspended in 1mL lysis buffer (deionized H_2_O with 50mM TrisHCl, 2mM DTT, 2 mM EDTA, 10% glycerol, 1% Triton). Samples were shaken with glass beads for 5 min, then combined in a 1:1 ratio with Nano-Glo® Luciferase solution (Promega, #N1150). A Promega GloMax plate reader was used to measure luminescence. Pooled samples were used because previous studies have demonstrated that mice within each cage are equally infected.

### T cell transfers

For T cell transfers, OT-I (CD45.1.2 background) and Smarta mice (CD45.1.1 background) were bred in house. To isolate OT-I CD8^+^ or Smarta CD4^+^ T cells, lymph nodes and spleens were harvested and leukocytes were obtained by mechanical dissociation over a 70 µm filter. Red blood cells were lysed by incubation for 5 min at room temperature in 1 mL of lysis buffer (0.864% ammonium chloride (Sigma-Aldrich) diluted in sterile deionized H_2_O), and then washed with complete RPMI (RPMI, 10% fetal calf serum, 0.1% beta-2-mercaptoethanol, 1% non-essential amino acids, 1% sodium pyruvate, and 1% pen-strep). OT-I and Smarta T cells were enriched by magnetic activated cell sorting (MACS) using the EasySep^TM^ Mouse CD8+ Isolation Kit or (StemCell #19853) or CD4+ T Cell Isolation Kit (StemCell #19852). OT-I and Smarta purities were verified (∼80-95%) using flow cytometry for TCR Vα2 (shared) and Vβ5.1/5.2 (OT-I) or Vβ8.3 (Smarta) expression. In some experiments, isolated T cells were incubated with CellTrace Violet dye (Invitrogen #C34557) for 20 minutes at 37°C prior to transfer. 1×10^4^-10^6^ OT-I T cells and 2×10^4^-1×10^6^ Smarta T cells were transferred by retroorbital injection into recipient mice.

### *In Vitro* Co-Culture

mLN were collected from 3-5 B6 or *Irf8+32*^-/-^ mice, pooled, and mechanically processed into a single cell suspensions over a 70 µm filter. DCs were then counted and enriched with the EasySep mouse pan-DC enrichment kit (StemCell #19863). DCs were incubated in 1 µM gp61 peptide (Genscript) for 20 minutes at 37°C, except for an aliquot to be used in an unstimulated control. DCs were then washed and resuspended in complete RPMI (recipe described above) at 200 cells/µL. The spleen was collected from a Smarta mouse and processed over a 70 µM filter. Smarta CD4^+^ T cells were isolated with the EasySep CD4^+^ T cell enrichment kit (StemCell #19852), labelled with CellTrace Violet (Invitrogen #C34557) and resuspended at 800 cells/µL in either complete RPMI or complete RPMI with 10 nM exogenous retinoic acid (RA) (Sigma #R2625) added. 100 µL of DC solution and 100 µL of Smarta solution were added to each well in triplicate of each condition, for a final 1:4 DC:T cell ratio. Co-cultures incubated at 37°C for 84-96 hours. Cells were collected and stained for flow cytometric analysis as described below.

### Integrin Blockade

For ⍺4β7 blocking experiments, infected WT C57BL/6J mice were treated intraperitoneally with 1 mg anti-⍺4β7 antibody (clone DATK32, BioXCell #BE0034) or 1 mg rat IgG2a isotype control (clone 2A3, BioXCell #BE0089) every 4 days starting 1 day prior to infection until the earlier between 12 dpi or time of sacrifice. The anti-⍺4β7 clone DATK32 binds to a combinatorial epitope of the integrin dimer and has been demonstrated to prevent its interaction with MAdCAM-1 (23, 45). Adequate binding of DATK32 to ⍺4β7 in our hands was demonstrated by decreased binding of a ⍺4β7 detection antibody of the same clone (Fig. 6A, B). For ⍺L blocking experiments, infected WT C57BL/6J mice were injected intraperitoneally with 650 ug anti-⍺L antibody (clone M17/4, BioXCell #BE0006) or IgG2a isotype control with the same regimen as ⍺4β7 blockade.

### Flow cytometry

Single-cell suspensions were prepared from intestinal sections by shaking diced tissue at 37°C for 20-30 min in Hank’s Balanced Salt Solution with 5 mM EDTA and 1 mM DTT, which separates the epithelial cells from the lamina propria. The lamina propria segments were rinsed in plain RPMI to remove EDTA, minced with scissors, and digested at 37°C for 30 min in RPMI with 0.16 mg/mL Liberase TL (Roche #5401020001) and 0.5 mg/mL DNase I (Roche #10104159001). Both the intestinal epithelial layer and lamina propria cells were centrifuged at 500 x g for 5 min, resuspended, then passed through 70 µm. This step was repeated with 40 µm filters. Ileal-draining mesenteric lymph nodes were harvested and dissociated through 70 µm filters. mLN were resuspended in cRPMI. Cells from all tissues were washed in FACS buffer (1x PBS, 0.2% bovine serum antigen, 1 mM EDTA), and incubated in Fc block (99.5% FACS Buffer, 0.5% normal rat IgG, 1 µg/ml 2.4G2) at 4°C for 15 min prior to staining. Cells were stained for cell death using Ghost Dye Violet 510 Viability Dye (Cytek #13-0870-T100) or Ghost Dye Red 780 (for experiments with CellTrace Violet, Cytek #13-0865-T100) in 1x PBS at 4°C for 15 min. Cells were washed after cell death staining and surface antibodies were added and stained at 4°C for 20-30 min. Cells were then washed in FACS buffer prior to acquisition. Cells were stained using the following fluorochrome-conjugated anti-mouse antibodies: BUV395 itg⍺V (clone RMV7, BD #747834), BUV496 CD4 (clone GK1.5, BD #612952), BUV563 CD8a (clone 53-6.7, BD #784535), BUV615 TCR v⍺2 (clone B20.1, BD #751416), BUV805 integrin ⍺L (clone 2D7, BD #741919), FITC integrin ⍺2 (clone DX5, BD #553857), FITC TCR Vβ5.1/2 (clone MR9-4, BD #553189) Percp-Cy5.5 integrin β7 (clone FIB27, BD #121008), RB780 integrin ⍺4 (clone R1-2, BD #755682), BV605 integrin ⍺E (clone 2E7, BioLegend #121433), BV650 Epcam (clone G8.8, BioLegend #118241), BV711 CD19 (clone 6D5, BioLegend #115555), BV711 NK1.1 (clone PK136, BioLegend #108745), ef450 CD45.1 (clone A20, eBioscience # 48-0453-82), APC CD45.1 (clone A20, BioLegend #110714), APC CD62L (clone MEL-14, BioLegend #104412), AF700 CD90.2 (clone 30-H12, BioLegend #105320), BV785 CD44 (clone IM7, BioLegend #103041), APC-Cy7 CD44 (clone IM7, BioLegend #103028), PE TCR Vβ8.3 (clone 1B3.3, BioLegend #156304), PE ⍺4β7 (clone DATK32, eBioscience #12-5887-83), Percp-ef710 ⍺4β7 (clone DATK32, Invitrogen #46-5887-82), PE integrin β1 (clone HMβ1.1, BioLegend #102207), PE-Cy7 integrin ⍺1 (clone HM⍺1, BioLegend #142608). Data were collected on an LSRFortessa or FACSymphony A3 Lite (BD Biosciences) and analyzed with FlowJo v10 software (TreeStar).

### Histopathology and Immunohistochemistry

To prepare intestinal sections mice were euthanized and the ileum removed, flushed with PBS, cut open longitudinally and ‘Swiss-rolled’. Samples were fixed overnight at 4°C in 10% neutral buffered formalin (Thermo Fisher) followed by 30% sucrose overnight (Thermo Fisher Scientific #15-188-51). Swiss rolls were then dehydrated and embedded in paraffin and 5 µm sections were cut and mounted on slides. For histopathologic analysis by hematoxylin and eosin staining, a Gemini Autostainer (Thermo Fisher Scientific) was used to deparaffinize the sections in xylene before rehydrating them by sequential baths in 100% ethanol, 95% ethanol and 70% ethanol followed by running water for 2 minutes. Hematoxylin and eosin stains were applied as previously published (46). Slides were imaged at 20X using the color camera mounted on a Leica DM6000 Widefield microscope.

Chromogenic IHC was performed as described elsewhere (47) using a Leica BOND RXm automated platform combined with the Bond Polymer Refine Detection kit (Leica #DS9800). Briefly, deparaffinized and rehydrated sections were pretreated with the epitope retrieval BOND ER2 high pH buffer (Leica #AR9640) for 20 min at 98°C. Endogenous peroxidase was inactivated with 3% H2O2 for 10 min at room temperature. Nonspecific tissue–antibody interactions were blocked by incubating the sections for 30 min at room temperature with Leica PowerVision IHC/ISH Super Blocking solution (PV6122). The same blocking solution also served as a diluent for the primary antibodies. Sections were incubated in primary antibody (clone MECA-367, Invitrogen #16-5997-85) for 45 min at room temperature. A biotin-free, polymeric detection system consisting of horseradish peroxidase conjugated anti-rabbit (Leica #DS9800), anti-rat (Vector Laboratories #MP7444), or anti-mouse (Leica #PV6114) IgG was then applied for 25 min at room temperature. Immunoreactivity was then revealed with the diaminobenzidine chromogen reaction. Tissue sections were finally counterstained in hematoxylin, dehydrated in an ethanol series, cleared in xylene, and permanently mounted with a resinous mounting medium (Thermo Scientific ClearVue coverslipper). Slides were imaged at 20X using the color camera mounted on a Leica DM6000 Widefield microscope.

### Bioinformatic Analysis

Following 10 days of *C. parvum* infection, single cell suspensions of the ileal draining lymph node, ileal lamina propria, and ileal epithelial cell layer were generated as described above and reported previously (48). Viable cells were used for CITE-seq profiling of mRNA and antibody-derived tag (ADT) protein expression (Biolegend catalog #199901). CITE-seq data were aligned to the mouse reference genome (GRCm39) using CellRanger (v8.0.0). Downstream analysis was performed in R (v4.5.1) primarily using the Seurat (v5.3.1) framework where cells were filtered for quality: RNA and ADT library sizes were restricted to ±2.5 median absolute deviations and cells containing >750 detected genes and <20% mitochondrial reads in the ileum. Newly generated samples were merged with existing GutPath intestinal CITE-seq dataset (PMID: 41497582; GEO: GSE313636) before normalization, integration and clustering. Harmony (v.1.2.4) integration was employed (via Seurat v5.3.1) to integrate newly generated data and the publicly available reference dataset (GutPath.org). Integration was performed separately for both the RNA and ADT modalities. Following integration, Weighted Nearest Neighbor (WNN) analysis was employed to construct a joint multi-modal neighbor graph used for UMAP dimensional reduction and clustering.

SingleR (v2.12.0) was used to assign cell type identities, utilizing the Gutpath.org reference (PMID: 41497582) to first establish broad cell lineages followed by iterative, subset-specific label transfer for more granular cell type annotations. This iterative process allowed for the high-resolution mapping of complex populations, including specific stromal subtypes, while maintaining fidelity to the published nomenclature. Heatmaps were constructed (ggplot2 v4.0.1) and used to visualize expression patterns of adhesion molecules (*Madcam1*, *Vcam1*, and *Icam1*). Statistical differences in expression across infection groups and conditions were evaluated using Wilcoxon rank-sum tests ggpubr (v0.6.2) within target cell lineages.

### Statistics

Statistical significance was calculated using unpaired t test with Welch’s correction to compare 2 groups or 2-way ANOVA followed by multiple comparisons when comparing 3 or more groups (see the grouped analyses in Figure 5). Statistical significance was calculated for area under the curve analyses using paired t tests. Significance level was set at 0.05. Statistical analyses for non-CITE-seq datasets were performed using GraphPad Prism v9 and 10.

### Data Availability

The authors declare that data supporting the findings of this study are available within the paper and its supplementary files. Naïve, *Cryptosporidium parvum* day 4, and *Yersinia pseudotuberculosis* ileum CITE-seq data are part of the GutPath intestinal CITE-seq dataset available at GEO: GSE313636. *Cryptosporidium* day 10 ileum CITE-seq data was generated for this study and has been submitted to GEO/SRA and is pending acceptance. Any additional data is available upon request by emailing the corresponding author.

## Results

### *Cryptosporidium-*specific T cells express diverse integrin profile distinct to priming phase of the response

To define the pattern of integrin expression by parasite-specific T cells during *Cryptosporidium* infection, 10000 CD8^+^ OT-I and 20000 CD4^+^ Smarta cells were co-transferred into C57BL6/J mice that were then infected with maCp-ova-gp61. In the absence of model antigens, transferred cells do not expand and are not detected in the mucosal tissues, but infection with maCp-ova-gp61 results in T cell expansion in the mLN and detectable populations in the intestinal epithelial layer (IEL) and lamina propria (LP) (41). Mice were sacrificed 7 days post-infection (dpi), a timepoint at which parasite-specific T cells are still being primed in the mLN and activated T cells have reached the gut. Cells were isolated from the IEL, LP, and mLN for analysis by flow cytometry as described in Methods. The parasite-specific OT-I and Smarta cells were identified based on their expression of the congenic marker CD45.1 along with TCR chain V⍺2. A panel of integrins associated with T cell trafficking (⍺4, ⍺1, ⍺2, ⍺V, ⍺L, ⍺E, β7, β1, and β2) was assessed for expression changes during infection. For this analysis, the OT-I and Smarta cells were compared to endogenous naïve (as defined by CD44 low expression) CD8^+^ and CD4^+^ T cells. In the mLN, the site of priming, Smarta cells showed no increase in ⍺V and ⍺1 expression over naïve cells but did have increased expression of ⍺2, ⍺4, ⍺L, ⍺E, β1, and β7 (Fig. 1A, B). CD44^lo^CD8^+^ T cells and OT-I T cells both constitutively express high levels of ⍺E, and while OT-I express more ⍺1 than the naïve CD8^+^ T cells, both populations express this integrin at a low constitutive level (Fig. 1C). Additionally, OT-I had significantly increased expression of ⍺2, ⍺4, ⍺L, β1, and β7 compared to naïve CD8+ T cells (Fig. 1C, D). Moreover, even when compared against endogenous CD44^hi^ T cells (the majority of intestinal T cells (49)), parasite-specific Smarta and OT-I cells expressed significantly higher levels of ⍺4, β7, ⍺L, and ⍺2 (Supp. Fig. 1). OT-I cells also expressed more β1 than endogenous CD44^hi^ comparators, while the Smarta expression of β1 as well as ⍺E is similar to this population (Supp. Fig. 1). Comparisons between transgenic cells and CD44^hi^ endogenous cells are shown only for the integrins that were differentially expressed in Figure 1.

**Figure 1:**
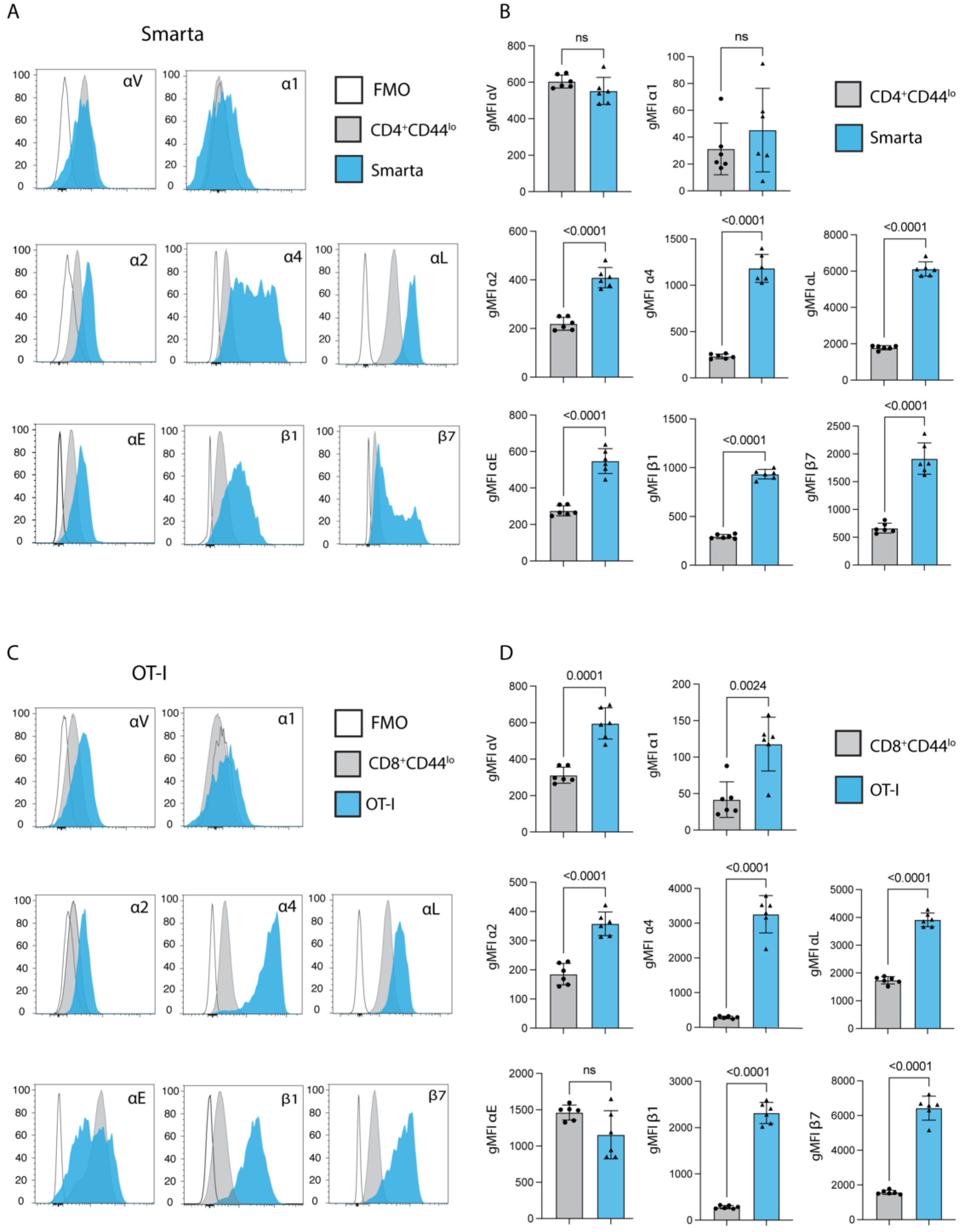
20000 Smarta cellCD19s and 10000 OT-I cells were transferred into wild type mice 1 days prior to infection with 50000 oocysts of maCp-ova-gp61. Expression of a panel of integrins in the mesenteric lymph node (mLN) 7 days post-infection (dpi) was measured by flow cytometry. **(A)** Histograms show fluorescence of these integrins on 2 cell populations in the mLN: endogenous naïve CD4^+^ T cells (Gating: Singlets>Live>CD19^-^NK1.1^-^CD90.2^+^>CD4^+^CD8^-^>CD45.1^-^CD44^lo^) and the Smarta T cells (Gating: Singlets>Live>CD19^-^NK1.1^-^CD90.2^+^>CD4^+^CD8^-^>V⍺2^+^CD45.1^+^). In each plot, these populations are concatenated across all mice and compared to CD4^+^ T cells from an mLN sample stained with the panel minus the integrin of interest (fluorescence minus one (FMO)). **(B)** Quantification of the comparison in A: Mean fluorescence intensity (gMFI) of every integrin in the panel for the endogenous naïve populations and the Smarta population in individual mice. **(C)** Histograms of fluorescence for the panel of integrins on 2 cell populations in the mLN: endogenous naïve CD8^+^ T cells (Gating: Singlets>Live>CD19^-^NK1.1^-^CD90.2^+^>CD4^-^CD8^+^>CD45.1^-^CD44^lo^) and adoptively transferred OT-I T cells (Gating: Singlets>Live>CD19^-^NK1.1^-^CD90.2^+^>CD4^-^CD8^+^> V⍺2^+^CD45.1^+^). These populations are concatenated across all mice and compared to CD8^+^ T cells from a mLN sample stained with an FMO for each integrin. **(D)** Quantification of the comparison in C: gMFI of every integrin in the panel for the endogenous naïve populations and the Smarta population in individual mice. Results representative of 3 independent experiments with n=6 mice/group. Bar plots show mean and SEM. Statistical significance determined by Welch’s t test. ns=p>0.05.

### Adhesion Molecule Ligand Expression During Infection

ICAM-1, VCAM-1, and MAdCAM-1 are ligands expressed in the stromal compartment which bind integrins to facilitate trafficking. To assess whether infection with *Cryptosporidium* alters gut expression of ICAM-1, VCAM-1, and MAdCAM-1, we generated new Cellular Indexing by Transcriptomes and Epitopes (CITE)-sequencing data and utilized a publicly available dataset from our group (48). This dataset contains gene expression data from the intestines of naïve mice and several enteric infections, including *Cryptosporidium parvum.* Expression from *Cryptosporidium*-infected mice (day 4 and 10) was compared with naïve mice and mice orally challenged with *Yersinia pseudotuberculosis* (day 5), which results in overt inflammation and significant disruption of the gut barrier (50) and is associated with high expression of these ligands on endothelial cells (Fig. 2A). Gene expression analysis revealed constitutive expression of *Icam1* and infection-induced expression of *Madcam1* and *Vcam1* on blood endothelial cells (Fig. 2A). Immunohistochemistry was used to assess MAdCAM-1 protein expression in the ileum of naïve and infected mice. There was cytoplasmic staining within cells of the lamina propria, but strong MAdCAM-1 staining was seen on endothelial cells of vessels within the gut-associated lymphoid tissues (GALT), particularly the Peyer’s patches, along the mucosal interface in both cohorts (red arrows, Fig. 2B). Morphologically, MAdCAM-1^+^ vessels within infected mice were characterized by diffusely plump endothelial cells compared to uninfected mice (Fig. 2B).

**Figure 2:**
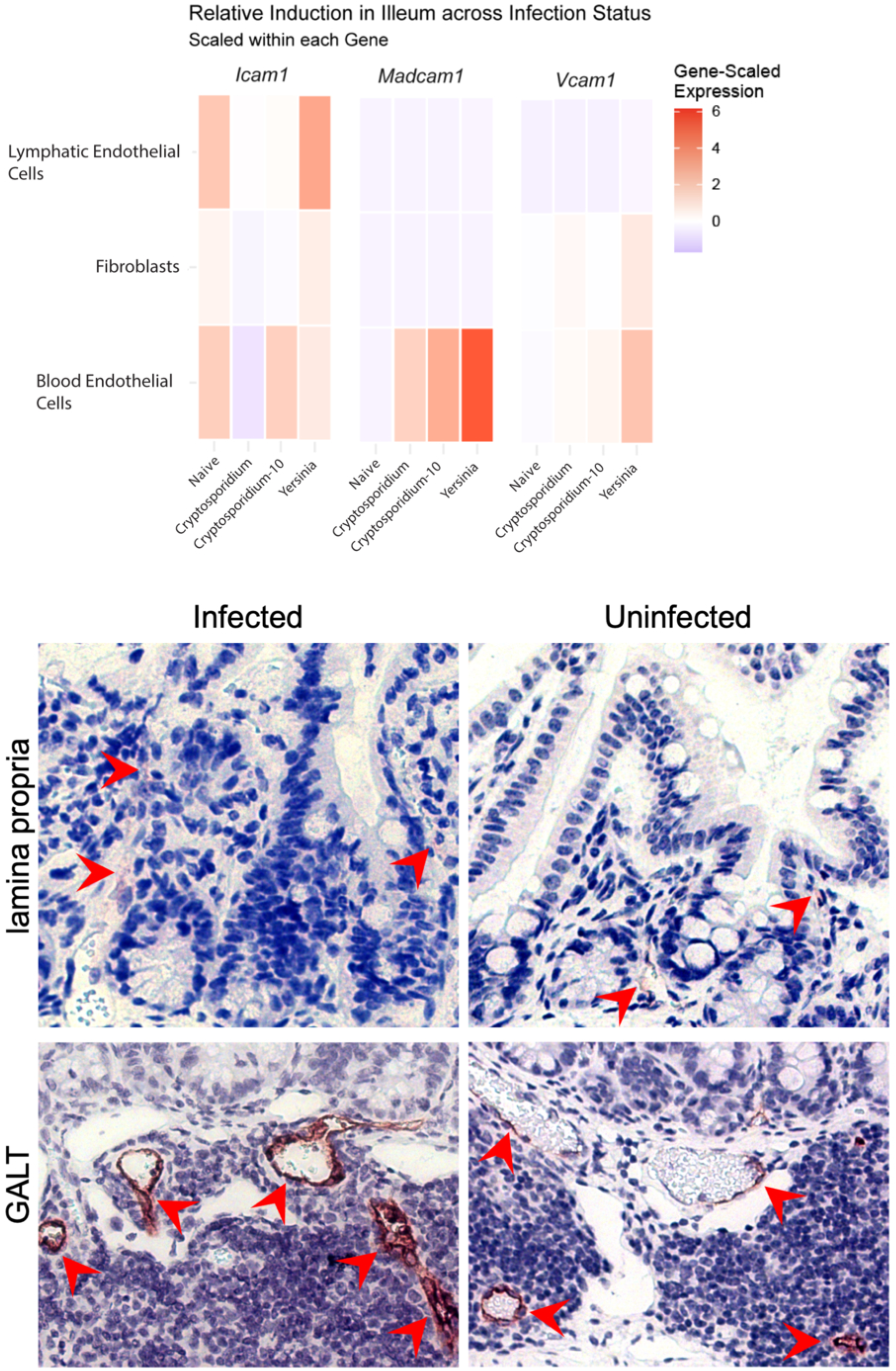
**(A)** Gene-scaled expression of *Icam1, Vcam1, and Madcam-1* from CITE-sequencing of the ileum (IEL and LP) of mice that are naïve, infected with *C. parvum* (4 and 10dpi), or infected with *Yersinia pseudotuberculosis* (5 dpi). n=1-2 mice/group. **(B)** Immunohistochemistry of MAdCAM-1 antigen localization within intestines of *Cryptosporidium* infected (7dpi) and uninfected mice. Arrows indicate MAdCAM-1^+^ endothelium in the GALT. n=3 mice/group.

### Integrin expression of *Cryptosporidium-*specific T cells is less distinct from endogenous CD44^lo^ population in the effector site

Among T cell populations in the small intestine, CD4^+^ T cells are more abundant in the LP and CD8^+^ T cells in the IEL (data not shown); therefore, these compartments were analyzed for Smarta and OT-I, respectively, on day 7 post-infection. Most intestinal T cells are CD44 intermediate or high during infection, but ∼10% of the endogenous CD4^+^ T cells in the LP and 20-30% of the endogenous CD8^+^ T cells in the IEL are CD44^lo^ and were used as a comparator for integrin expression. While still significantly elevated on the Smarta cells in the LP, expression of ⍺4, β7, and β1 is more similar between naïve and Smarta populations than was observed in the mLN, while Smarta ⍺L expression remained highly elevated over the naïve population (Fig. 3A, B). The same is true for the OT-I in the IEL, which also maintain high levels of ⍺E similar to the endogenous naive cells (Fig. 3C, D). At 60 dpi, a memory timepoint when infection has cleared, both Smarta and OT-I cells still expressed ⍺L, but Smarta cells in the gut had downregulated ⍺E compared to 10 dpi, while the OT-I memory population remained high for ⍺E (Supplemental Fig. 1E). Thus, by the time parasite-specific T cells have trafficked to the ileum, they are not as distinct from naïve populations in their integrin expression profile as was seen in the mLN.

**Figure 3:**
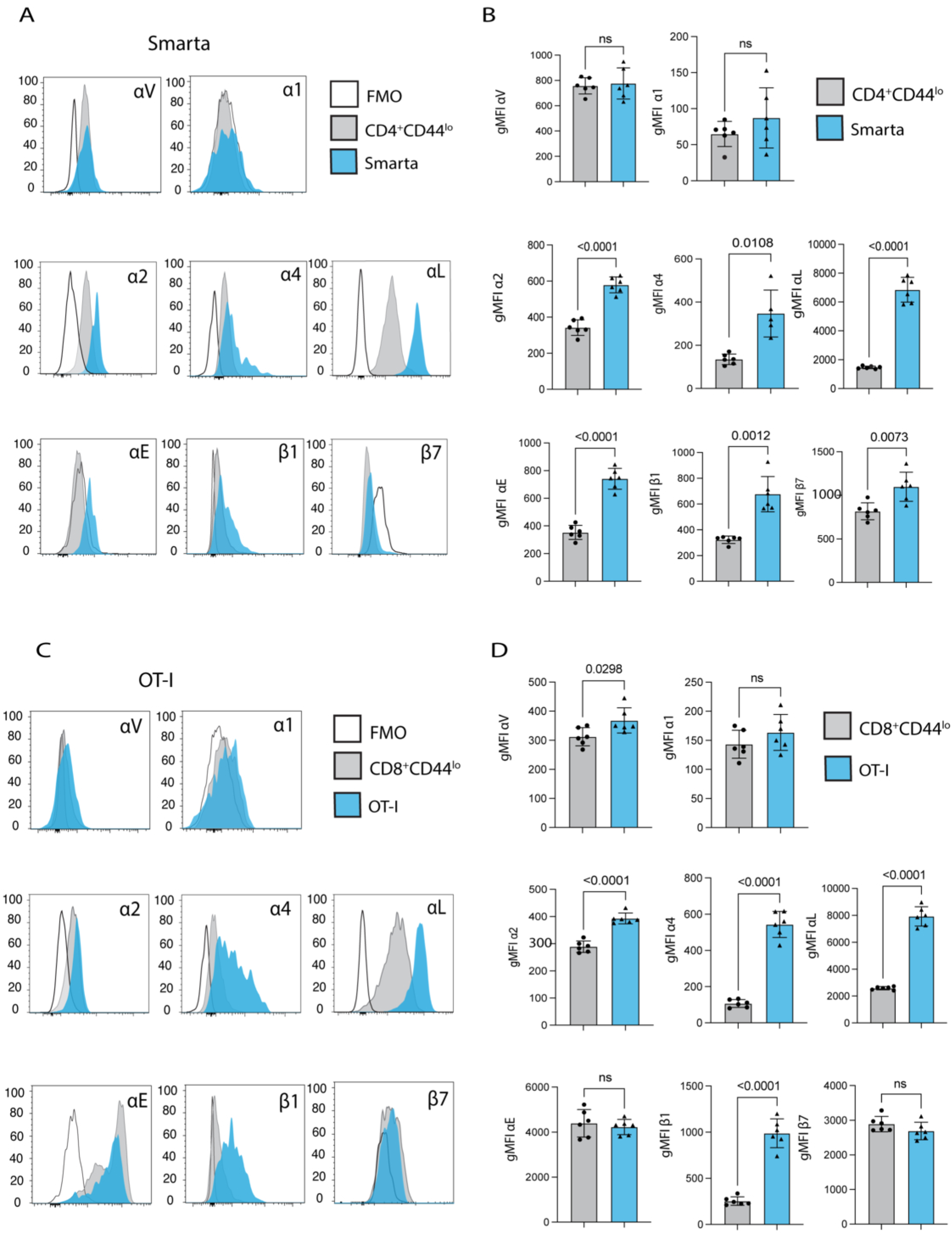
20000 Smarta cells and 10000 OT-I cells were transferred into wild type mice 1 day prior to infection with 50000 oocysts of maCp-ova-gp61. Expression of a panel of integrins in the gut 7dpi was measured by flow cytometry. **(A)** Histograms of fluorescence for a panel of integrins on 2 cell populations in the ileum lamina propria (LP) 7 dpi: endogenous naïve CD4^+^ T cells (Gating: Singlets>Live>CD19^-^NK1.1^-^CD90.2^+^>CD4^+^CD8^-^>CD45.1^-^CD44^lo^) and adoptively transferred Smarta T cells (Gating: Singlets>Live>CD19^-^NK1.1^-^CD90.2^+^>CD4^+^CD8^-^ >V⍺2^+^CD45.1^+^). In each plot, these populations are concatenated across all mice and compared to the T cells from an intestinal epithelial layer (IEL) sample stained with the panel minus the integrin of interest (FMO). **(B)** Quantification of the comparison in A: gMFI of every integrin in the panel for the endogenous naïve populations and the Smarta population in individual mice. **(C)** Histograms of flow cytometry fluorescence for the panel of integrins on 2 cell populations in the mLN 7 dpi: endogenous naïve CD8^+^ T cells (Gating: Singlets>Live>CD19^-^NK1.1^-^CD90.2^+^>CD4^-^CD8^+^>CD44^lo^CD45.1^-^) and adoptively transferred OT-I T cells (Gating: Singlets>Live>CD19^-^NK1.1^-^CD90.2^+^>CD4^-^CD8^+^>Va2^+^CD45.1^+^). In each plot, these populations are concatenated across all mice compared to the T cells from an IEL sample stained with the panel minus the integrin of interest (FMO). **(D)** Quantification of the comparison in C: gMFI of every integrin in the panel for the endogenous naïve populations and the Smarta population in individual mice. Results representative of 3 independent experiments with n=6 mice/group. Bar plots show mean and SEM. Statistical significance determined by Welch’s t test. ns=p>0.05.

### Impact of cDC1 and RA on the *Cryptosporidium*-specific T cell integrin expression

At homeostasis, CD103^+^ cDC1s constitute the major source of RA in the mLN, which promotes expression of ⍺4β7 on newly activated T cells (51). Our previous studies have shown that in the absence of cDC1s, Smarta T cells proliferate, but do not upregulate ⍺4β7 and do not traffic to the small intestine (52). To understand the contribution of cDC1s to the expression of integrins during infection, *Irf8+32^-/-^* mice that lack cDC1s (53) were utilized. Because in these mice OT-I T cells do not expand, only Smarta T cells were analyzed. Prior to transfer into WT or *Irf8+32^-/-^* mice, Smarta T cells were labelled with CellTrace Violet (CTV) to track expansion. Mice were subsequently infected with maCp-ova-gp61, and T cell expansion and expression of integrins was assessed at 7 dpi. In the *Irf8+32^-/-^* mice, Smarta cells expressed less ⍺4, β7, β1, and ⍺L than Smarta cells in WT mice (Fig 4A, B). The numbers of Smarta cells in the mLN indicated that they had expanded significantly less in the *Irf8+32^-/-^* compared to WT mice (data not shown and previously published (52)) and CTV dilution confirmed that the T cells experience reduced proliferation in the absence of cDC1s (Fig 4C). Nevertheless, when the cells in the latest round of division (i.e. the fully differentiated effector T cells) were analyzed, the expression of ⍺4, β7, and ⍺L remained significantly reduced, but the difference in β1 expression in the fully divided cells was mitigated. These data sets indicate that cDC1s are required for optimal expansion of Smarta T cells but are also linked to ⍺4β7 and ⍺L.

**Figure 4:**
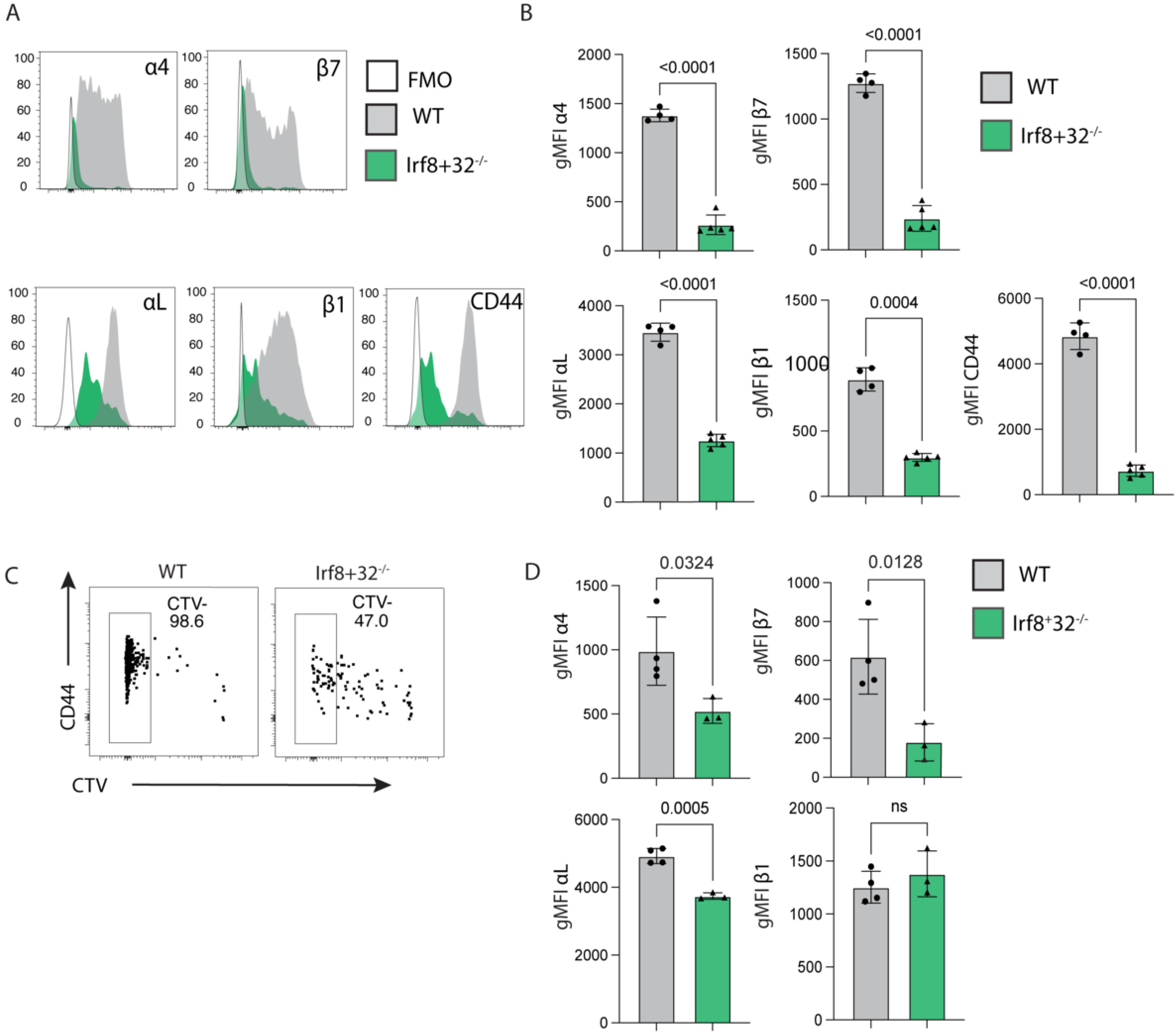
**(A and B)** 20000 Smarta T cells were transferred into either wild type mice or *Irf8+32^-/-^* mice 1 day prior to infection with 50000 oocysts. Integrin expression on the Smarta population was measured in the mLN 7 dpi by flow cytometry. **(A)** Histograms of fluorescence for a selection of differentially expressed integrins on Smarta cells. In each plot, these populations are concatenated across all mice and compared to a wild type mLN sample stained with the panel minus the integrin of interest (FMO). **(B)** Quantification of the comparison in A: gMFI of each integrin for the endogenous naïve populations and the Smarta population in individual mice. Panels A and B are representative of 4 independent experiments, n=3-5 mice/group. **(C and D)** 20000 Smarta T cells were labelled with CellTrace Violet (CTV) proliferation dye and transferred into either wild type mice or *Irf8+32^-/-^* mice 1 day prior to infection with 50000 oocysts. **(C)** Proliferation and activation of these cells was demonstrated by CTV dilution and CD44 expression. Cells concatenated across all mice, and a gate shows the most divided (CTV-) cells. **(D)** Expression of CD44 versus CTV staining of the concatenated Smarta cells, with a gate on the last 2 cell divisions (CTV^-^). gMFIs of ⍺4, β7, ⍺L, and β1 expression on the CTV^-^ population was compared between WT and *Irf8+32^-/-^* mice. Panels C and D are representative of 1 independent experiment, n=3-4 mice/group. Bar plots show mean and SEM. Statistical significance determined by Welch’s t test. ns=p>0.05.

To test whether cDC1-derived RA contributes to β7, β1, ⍺4, and ⍺L expression, DCs were isolated from the mLN of naïve WT or *Irf8+32^-/-^* mice, pulsed with the cognate peptide gp61-80, and co-cultured with Smarta cells for 96 hours with or without exogenous RA. In the setting of a controlled DC:T cell ratio, the Smartas expand to this same extent in the WT-DC and *Irf8*-DC groups as demonstrated by dilution of the CTV labelling (Fig. 5A). When the expression of ⍺4, β7, β1, and ⍺L on the T cells was assessed, the Smarta cells cultured with DCs from *Irf8+32^-/-^* mice express less ⍺4β7 compared to co-culture with DCs from WT mice, and the addition of exogenous RA rescues ⍺4β7 expression (Fig. 5B, C). However, the absence of cDC1s did not affect the expression of ⍺L (Fig. 5D) or β1 (Fig. 5E) *in vitro.* While overall expression of ⍺L was not increased by exogenous RA (Fig. 5D), RA modestly enhanced the expression of β1, particularly on T cells cultured with the I*rf8+32^-/-^* DCs (Fig. 5E). This result confirms the key role of RA in ⍺4 and β7 expression on T cells and reveals potential regulation of β1 by RA, while RA appears to have minimal effect on β1 expression.

**Figure 5:**
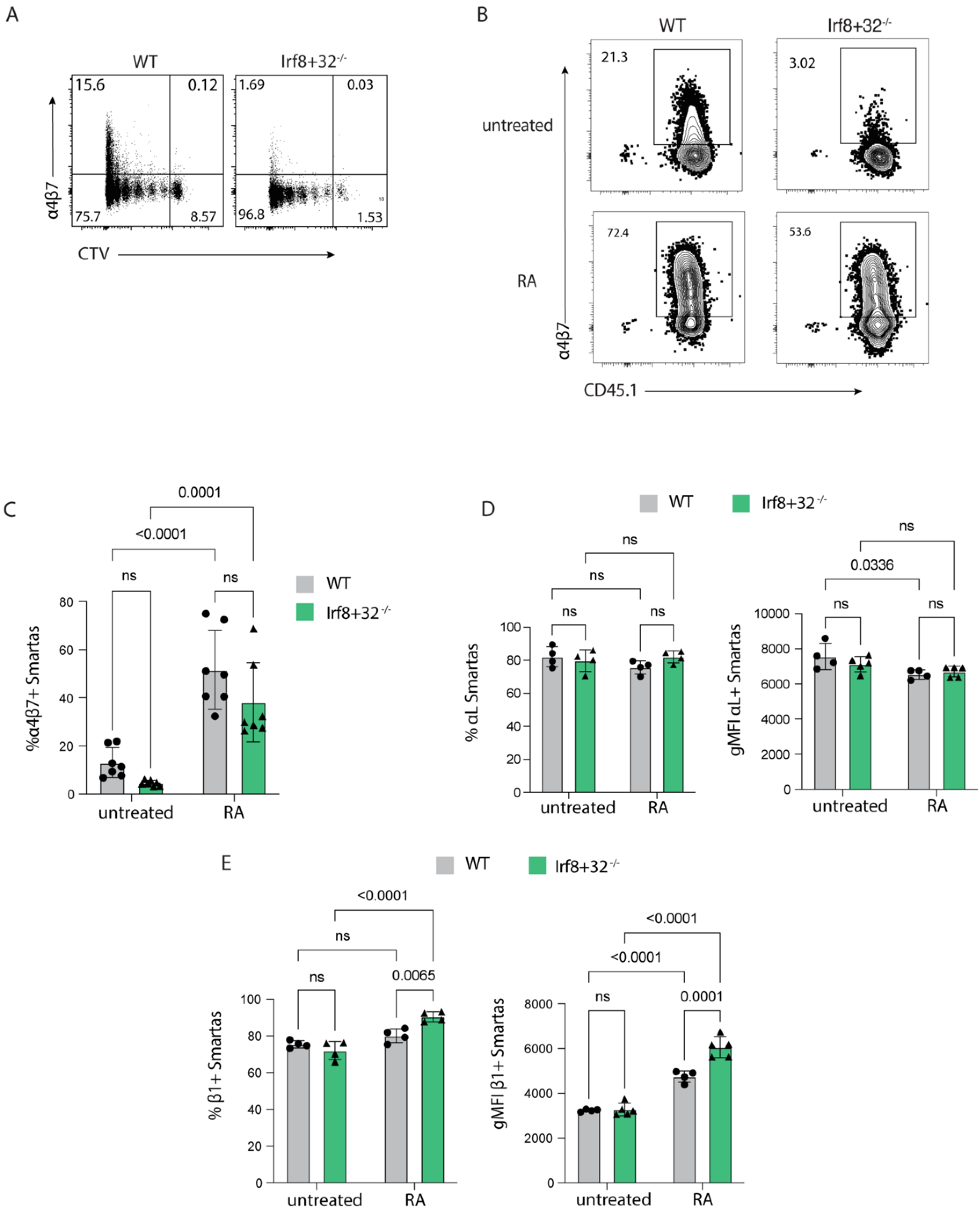
CellTrace Violet (CTV)-labelled Smarta T cells were co-cultured with DCs that were enriched from the mLN of WT or *Irf8+32^-/-^* mice and pulsed with gp61 peptide. Co-cultures incubated for 96h. Culture media was either untreated or with 10 nM exogenous RA. **(A)** Flow cytometry plots of CTV staining demonstrating the extent that the Smarta cells proliferation after co-culture. Representative wells from the untreated group. **(B)** Representative flow cytometry plots showing ⍺4 and β7 expression for the different DC sources (WT or *Irf8+32^-/-^)* with or without exogenous RA. **(C)** Quantification of ⍺4β7 expression on the groups shown in (B). **(D)** Quantification of ⍺L expression in the 4 conditions, shown as both positive frequency of Smarta and the gMFI of the positive Smarta cells. **(E)** Quantification of β1 expression in the 4 conditions, shown as both positive frequency of Smarta and the gMFI of the positive Smarta cells. Plots in **(C)** contain data from 2 independent experiments and plots in **(D)** and **(E)** show representative data from one of these experiments. Bar plots show mean and SEM. Statistical significance was determined by 2-way ANOVA with post-test multiple comparisons. ns=p>0.05.

### ⍺4β7 blockade does not prevent T cell trafficking to the ileum in *Cryptosporidium* infection

To evaluate the contribution of ⍺4β7 to T-cell-mediated control of *Cryptosporidium*, infected mice were treated with the monoclonal antibody DATK32 every 4 days for the duration of the experiment. Clone DATK32 is a well characterized reagent that blocks ⍺4β7 binding to its ligand MAdCAM-1 (21, 23, 45) and is routinely used *in vivo* (31, 54, 55). Consistent with these reports, *in vivo* treatment with DATK32 resulted in a reduced ability to detect ⍺4β7 on the mLN Smarta cells (Fig. 6A, B). Surprisingly, ⍺4β7 blockade did not affect *Cryptosporidium* burden in WT mice (Fig. 6C). In *Ifng*^-/-^ mice, which are highly susceptible to *Cryptosporidium parvum* and exhibit a CD4 T cell-dependent mechanism of resistance (9), blockade of ⍺4β7 did not increase parasite burden, as demonstrated by area under the curve comparisons from 4 independent parasite burden experiments in either WT or *Ifng*^-/-^ mice (Fig. 6D). Consistent with this result, when Smarta and OT-I cells were adoptively transferred into WT mice and the intestines were analyzed by flow cytometry 10 days post-infection, mice treated with anti-⍺4β7 exhibited similar OT-I and Smarta responses, in both frequency and number, in the IEL and LP compared with isotype controls (Fig. 6E, F).

**Figure 6:**
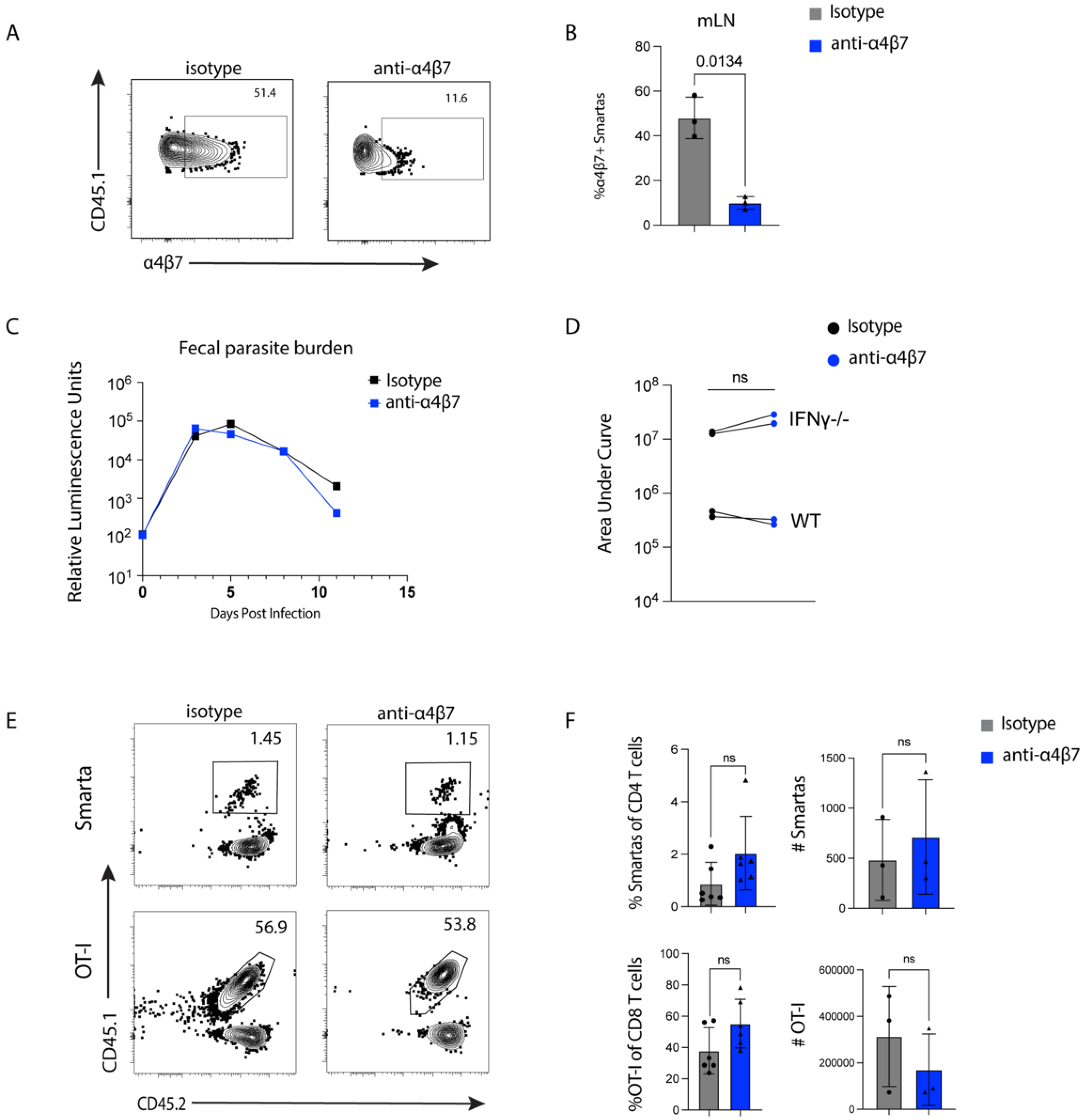
20000 Smarta cells and 10000 OT-I cells were transferred into wild type mice 1 day prior to infection with 50000 oocysts of maCP-ova-gp61. Mice were treated with 1mg each of either anti-⍺4β7 or isotype control 1 day prior to infection every 4 days until endpoint. **(A)** The blocking antibody’s binding to its target was validated through staining treated samples with a detection antibody of the same clone. Flow cytometry plots of ⍺4β7 detection on Smarta cells in the mLN of representative mice treated with either blocking antibody or isotype control on 10 dpi. **(B)** Quantification of the data in (A). **(C)** Parasite burden over time assayed by relative luminescence of the pooled feces from mice treated with either isotype or blocking antibody. n=3 mice/group. **(D)** The areas under the curve (AUC) from 4 independent measurements of fecal parasite burden after isotype or blocking antibody. 2 of the experiments were conducted in IFNɣ-/- mice as noted. n=3 mice/group. **(E)** Representative flow cytometry plots on 10 dpi showing Smarta and OT-I T cell frequencies in the guts of mice treated with either isotype control or blocking antibody. Smarta cells are shown in the LP and OT-I cells are shown in the IEL. **(F)** Quantification of the data in (E) as both frequency of parent populations and cell number. Representative of 3 independent experiments with n=3-4 mice/group. Statistical significance determined by Welch’s t test. ns=p>0.05.

### Blocking the interaction of integrin ⍺L with ICAM-1 inhibits T cell response to Cryptosporidium

Integrin ⍺L (CD11a) is upregulated on parasite-specific T cells compared to polyclonal CD44^lo^ controls (Fig. 1) and its ligand ICAM-1 is constitutively expressed on endothelial cells in the gut (Fig. 2A). This integrin has several known functions in the activation (56–58) and trafficking (36, 59) of effector T cells. Indeed, one of the functions of ⍺Lβ2 integrin is to facilitate entry of lymphocytes into lymph nodes, so the effect of ⍺L blockade on seeding of the Smarta cells into the mLN was assessed. 1×10^6^ cells were transferred concurrently with isotype or anti-⍺L treatment, and the Smarta numbers in the mLN and spleen were analyzed 24 hours later. There were seven-fold fewer Smarta cells in the mLN of anti-⍺L treated mice consistent with impaired seeding of LNs, but the antibody blockade did not affect numbers of Smarta cells in the spleen (data not shown).

To assess the effect of ⍺L blockade on Smarta T cell priming, 1×10^6^ cells were labelled with CTV before adoptive transfer into mice that were treated with either isotype or blocking antibody, and Smarta priming was assessed in the mLN at 7dpi. CTV staining revealed that ⍺L blockade at the same time as T cell transfer did not affect T cell priming (Fig 7A). This experiment was repeated with antibody treatment starting 48h post-T cell transfer. Again, the Smarta cells in the mLN of treated mice underwent similar proliferation and upregulation of activation markers compared to the isotype control (Fig. 7B). The defect in Smarta number in the mLN was mitigated by the delayed initiation of blockade which allowed for seeding of the LN by naïve Smarta T cells (Fig. 7B), so subsequent experiments were done with this regimen.

**Figure 7:**
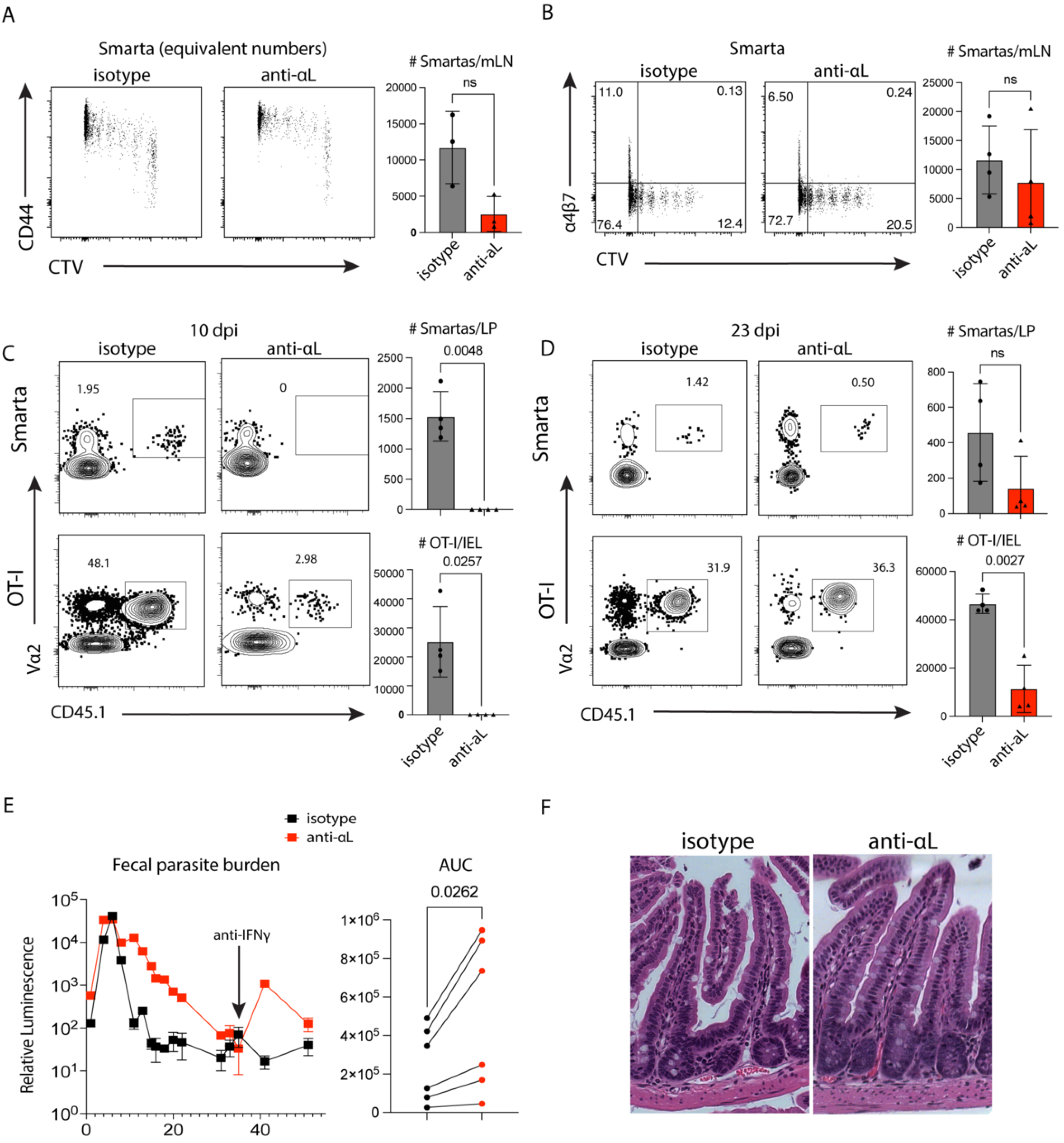
1×10^6^ CellTrace Violet labelled Smartas were transferred into WT mice 1 day prior to infection with 50000 maCp-ova-gp61. Treatment with 650 µg isotype control or anti-⍺L was given 1 day prior to infection in **(A)** and 2 days post-infection in **(B)**. **(A)** CD44 expression versus CTV dilution of the concatenated Smarta cells across all mLN at 7dpi, downsampled to show equal number for both groups. Bar plot quantification of Smarta number per mLN. Representative of 2 independent experiments, n=3 mice/group. **(B)** Integrin ⍺4β7 expression against CTV staining on Smarta cells in the mLN 7 dpi, followed by bar plot quantification of Smarta number per mLN. Data from 1 independent experiment, n=4 mice/group. **(C and D)** 20000 Smarta cells and 10000 OT-I cells were transferred into WT mice 1 day prior to infection with 50000 oocysts of maCP-ova-gp61. Treatment with 650 µg isotype control or anti-⍺L was started 2 dpi and given again on d6 and d10. **(C)** Representative flow cytometry plots showing effector T cell frequencies at 10 dpi in the ileum of mice treated with either isotype control or blocking antibody. Bar plot quantification showing the Smarta and OT-I number per sample. Representative of 2 independent experiments, n=3-4 mice/group. **(D)** Representative flow cytometry plots showing effector T cell frequencies at 23 dpi in the ileum from representative mice treated with either isotype control or blocking antibody. 1 independent experiment, n=4 mice/group. Bar plots are quantification of the Smarta and OT-I responses in (D) as absolute number per sample. **(E)** Parasite burden over time assayed by relative luminescence of the pooled feces from mice treated with either isotype or blocking antibody. A bolus of anti-IFNɣ was given at 31 dpi to assess clearance. Adjacent plot shows the areas under the curve (AUC) from 5 independent measurements of fecal parasite burden kinetics. **(F)** Representative images of H&E staining of sections from fixed ileal Swiss rolls from mice treated with either isotype control or blocking antibody. ns=p>0.05.

When the effect of ⍺L blockade on trafficking to the gut was examined 10 days post-infection, there was a significant decrease in parasite-specific T cells in the intestine after ⍺L blockade (Fig. 7C). However, when Smarta and OT-I numbers in the intestine were analyzed 23 days post-infection, it was found that after ⍺L blockade, cells eventually trafficked to the ileum, but numbers remained lower than in the isotype control mice (Fig. 7D). Nevertheless, blocking ⍺L and the associated T cell deficit led to higher fecal *Cryptosporidium* burden starting 6 days post-infection and a failure to fully clear the infection, as evidenced by parasite burden recrudescence after depletion of IFNɣ at 31 dpi (Fig. 7E). Hematoxylin and eosin staining of Swiss-rolled ileum sections revealed that the anti-⍺L treated mice exhibited increased crypt length and branching, a pathology associated with high parasite burden (Fig 7F) (60). These data support that ⍺L blockade leads to a defect in T-cell-mediated control.

## Discussion

The location of effector T cells is a key aspect of the immune response to pathogens that have highly specific tissue tropism, and targeting cells to these sites is a challenge for vaccine design. An improved understanding of the mechanisms by which T cells traffic to the intestine during *Cryptosporidium* infection sheds light on pathways that a potential vaccine could leverage to generate memory T cells in the gut. *Cryptosporidium* as an acute infection model is unique in its restriction to epithelial cells of the small intestine, serving as an experimental system to understand highly localized ileal inflammation. Here we have used a combination of transgenic parasites and TCR transgenics to better understand how parasite-induced T cell activation influences integrin expression and how this is altered as these cells access the gut. We also assessed how integrin blockade affects T cell trafficking and control of this enteric infection. Compared to previous studies of colitogenic T cells (61), this study focused on the mechanisms of trafficking specifically to the distal small intestine in the setting of local infection associated with a T_h_1 T cell response. This survey demonstrated that activated parasite-specific T cells express a variety of integrins that distinguish them from other T cells in the mLN and gut.

The ability of cDC1s to promote resistance to *Cryptosporidium* is most closely linked to their ability to present antigen and produce IL-12, but the identification of high levels of the canonical gut-homing integrin ⍺4β7 on parasite-specific T cells in the mLN suggests that cDC1 production of RA is another important function, as has been reported in other models (51). In addition, our datasets reveal that cDC1s are required for maximal ⍺L and β1 expression on parasite-specific T cells. While it is known that RA signaling activates transcription of the β7 integrin, exogenous RA did not greatly affect ⍺L and β1 expression on T cells *in vitro.* Thus, the loss of these other integrins in the absence of cDC1s is not easily explained by decreased RA signaling. Further studies are needed to evaluate RA-independent functions of cDC1s that may influence the homing behavior of these pathogen-specific T cells.

Previous reports that cDC1s are important for T cell trafficking during *Cryptosporidium* infection and promote ⍺4β7 expression (52) supported the hypothesis that this integrin would contribute to the ability of parasite-specific T cells to migrate to the site of infection. However, antibody blockade of ⍺4β7 did not affect Smarta and OT-I numbers in the gut or alter the host ability to control infection. Prior studies have revealed the requirement of ⍺4β7 for gut-homing to be context-dependent (62). For example, ⍺4β7 blockade affected T cell frequencies in a mouse model of chronic but not acute colitis (29). In addition, while there has been an emphasis on ⍺4β7 in gut homing (31, 63), other integrins such as ⍺Lβ2 and ⍺4β1 also contribute to this in IBD models (64, 65), but less is known about these integrins in an infectious context. Our observation of ⍺4β7-independent trafficking to the ileum during *Cryptosporidium* infection supports the existence of alternative gut-homing pathways. These findings exist in the clinical context of vedolizumab, a blocking antibody against the human ⍺4β7 heterodimer (66), which is used to treat inflammatory bowel disease (67). While this treatment has been used extensively in patients, it is not associated with an increased risk of cryptosporidiosis (68)—a common opportunistic infection in the immunocompromised host (5).

The integrin ⍺L, which dimerizes with β2, has a complex biology with demonstrated roles in T cell priming and trafficking. In a model of oral antigen delivery, even when T cells were pre-activated *in vitro* to control for their level of expansion and differentiation, adoptively transferred β2 ^-/-^ T cells exhibited a defect in trafficking to the lamina propria (36). Similarly, the studies presented here show that blockade of ⍺L was sufficient to delay the accumulation of *Cryptosporidium*-specific T cells in the small intestine. These results need to be interpreted with care, as it is important to consider that ⍺L has established roles in T cell activation. For example, the interaction of ⍺L with ICAM-1 has been shown to stabilize the TCR:MHC interaction (56), and in models of vaccination and infection ⍺L blockade can compromise T cell expansion (58, 69). As expected, pretreatment of mice with anti-⍺L reduced the ability of naïve Smarta T cells to seed the mesenteric lymph node but did not result in obvious differences in T cell priming as measured by proliferation and activation markers. Moreover, delayed treatment with the blocking antibody still resulted in delayed ability of T cell to traffic to the gut and control *Cryptosporidium*. While it is possible that ⍺L affects other aspects of T cell priming, our observations suggest no requirement for ⍺L in initial T cell activation during *Cryptosporidium* infection but rather a role in trafficking. Recognition of the contribution of ⍺L to gut-homing indicates that induction of this integrin could be a useful immunological benchmark in mucosal vaccine design.

Since the recognition of integrins and their ligands as a mechanism for tissue-specific lymphocyte trafficking, subsequent studies have added nuance to our knowledge of these pathways by analyzing their role in lymphocyte subsets and different inflammatory contexts. Several studies have demonstrated that the rules of tissue targeting established during homeostasis are less stringent during overt inflammation, when activating signals result in the increased expression of adhesion molecules on lymphocytes and peripheral endothelial cells. Our work, in the context of this literature and the clinical efficacy of ⍺4β7 blocking antibodies in IBD, is an important reminder that there is more to learn about the rules that govern mucosal T cell trafficking.

## Acknowledgements

The data for this (manuscript or presentation) were generated in the Penn Cytomics and Cell Sorting Shared Resource Laboratory at the University of Pennsylvania (RRID:SCR_022376). Penn Cytomics is partially supported by the Abramson Cancer Center NCI Grant (P30 016520). H&E staining and immunohistochemistry were performed by the Comparative Pathology Core (CPC) at PennVet and histological evaluations were performed by CPC staff (J. Engiles).

The Center for Host-Microbe Interactions (CHMI) at the University of Pennsylvania generated the GutPath CITE-seq dataset probed in Figure 2. GutPath is an endeavor funded by the NIH Mucosal Immunology Studies Team (MIST).

This work was supported in part by the National Institutes of Health with grants to C.A. Hunter and B. Striepen (R01AI148249), B. Striepen (R01AI112427), a fellowship and training grant support to M.I. Merolle (F30AI186259 and T32007632), training grant support to B.E. Haskins (T32AI007532), a fellowship to I.S. Cohn (F30AI169744), and training grant support to C.A. Howard (T32AI055428). B. Striepen and C.A. Hunter are supported by the Commonwealth of Pennsylvania.

## Author contributions

M.I. Merolle: Conceptualization, Data curation, Formal analysis, Investigation, Methodology, Resources, Software, Validation, Visualization, Writing—original draft, Writing—review & editing, B.E. Haskins: Conceptualization, Investigation, Methodology, Writing—review and editing, J. Engiles: Methodology, Formal analysis, writing-review and editing, A. Hart: Formal Analysis, Methodology, Writing—review and editing, I.S. Cohn: Investigation, Writing-review and editing, K.M. O’Dea: Investigation, C.A. Howard: Investigation, Writing—review & editing, D.A. Christian: Resources, Supervision, J.H. Byerly: Resources, B. Striepen: Conceptualization, Funding acquisition, Resources, Writing—review & editing, C.A. Hunter: Conceptualization, Funding acquisition, Methodology, Project administration, Supervision, Validation, Writing—original draft, Writing—review & editing.

---

**Supplemental Figure 1:** 20000 Smarta cells and 10000 OT-I cells were transferred into WT mice 1 day prior to infection with 50000 maCp-ova-gp61 oocysts. **(A)** Expression of select integrins in the mLN at 7dpi was measured by flow cytometry. The histograms show fluorescence of these integrins on 2 cell populations in the mLN: endogenous antigen-experienced (CD45.1^-^CD44^hi^) CD4^+^ T cells and the Smarta T cells. In each plot, these populations are concatenated across all mice and compared to CD4^+^ T cells from an mLN sample stained with the panel minus the integrin of interest (FMO). **(B)** Quantification of the comparison in A: gMFI of select integrins for the endogenous CD44^hi^ populations and the Smarta population in individual mice. **(C)** Histograms of fluorescence for the panel of integrins on 2 cell populations in the mLN: endogenous antigen-experienced (CD45.1^-^CD44^hi^) CD8^+^ T cells and adoptively transferred OT-I T cells. In each plot, these populations are concatenated across all mice and compared to CD8^+^ T cells from a mLN sample stained with the panel minus the integrin of interest (FMO). **(D)** Quantification of the comparison in C: gMFI of the selected integrins for the endogenous CD44^hi^ populations and the OT-I population in individual mice. Results representative of 3 independent experiments with n=5-6 mice/group. Bar plots show mean and SEM. Statistical significance determined by Welch’s t test. ns=p>0.05. **(F)** Flow cytometry plots showing select integrin expression on Smarta and OT-I cells in the LP and IEL at 10 dpi and 60 dpi, concatenated across 5 mice. Representative of 2 independent experiments.

